# Targeting LDHC dysregulates the cell cycle and improves sensitivity to cisplatin and olaparib

**DOI:** 10.1101/2021.03.03.433525

**Authors:** Adviti Naik, Julie Decock

## Abstract

The cancer testis antigen (CTA) lactate dehydrogenase C (LDHC) is a promising anti-cancer target with tumor-specific expression, immunogenicity and a role in metabolic reprogramming. Interrogation of the TCGA breast cancer cohort demonstrates upregulation of *LDHC* expression, conferring unfavorable prognosis. Although the role of LDHC is well characterized in spermatocytes, its role in tumors remains largely unknown. We investigated whether LDHC is involved in regulating genomic stability and may be targeted to affect tumor cellular fitness. Silencing *LDHC* in four breast cancer cell lines significantly increased the presence of giant cells and nuclear aberrations, DNA damage and apoptosis. *LDHC* silenced cells demonstrated aberrant cell cycle progression with differential expression of cell cycle checkpoint and DNA damage response regulators. In addition, *LDHC* silencing induced microtubule destabilization, culminating in increased mitotic catastrophe and reduced long-term survival. Notably, cisplatin and olaparib treatment further reduced survival of *LDHC* silenced cells. This study supports the therapeutic potential of targeting LDHC to mitigate cancer cell survival, and improve sensitivity to DNA damaging and DNA damage repair inhibiting agents.

## 1. INTRODUCTION

Breast cancer remains the most common cancer in women and has now overtaken lung cancer as the leading cause of cancer-related death in women (Bray *et al*, 2018). The clinical and molecular heterogeneity of breast cancer has led to the definition of numerous subtypes. The immunohistochemically-defined triple negative breast cancer (TNBC) subtype is an aggressive subgroup of breast tumors with high early recurrence rates and poor clinical outcome. Furthermore, TNBC patients do not benefit from targeted therapy due to the absence of estrogen receptor (ER), progesterone receptor (PR) and human epidermal growth factor receptor 2 (HER2) receptor expression. More recently, gene expression profiling identified intrinsic subtypes with improved prognostic stratification, which are independent of standard clinicopathological variables (Lehmann & Pietenpol, 2014; Perou *et al*, 2000). Among these subtypes, basal-like breast cancer is associated with the worst prognosis (Caan *et al*, 2014). Triple negative and basal-like breast cancer have a high degree of overlap with TNBCs accounting for about 80% of the latter and being associated with worse clinical outcome compared to non-basal-like TNBCs (Ahn *et al*, 2016; Liu *et al*, 2016; Parker *et al*, 2009; Seal & Chia, 2010). To date, numerous efforts have been made towards understanding the molecular underpinnings of basal-like breast cancer with the goal to identify novel and selective targets.

Cancer testis antigens (CTAs) constitute a group of proteins that are endogenously expressed in human germ cells and placental tissue, and are re-expressed in numerous cancer types (Gjerstorff *et al*, 2015). Lactate dehydrogenase C (LDHC) is a CTA that belongs to the lactate dehydrogenase (LDH) family of isozymes, comprising of LDHA, LDHB and LDHC (Mishra & Banerjee, 2019). LDHA and LDHB are the predominant LDH isozymes consisting of four LDH-M and LDH-H subunits respectively, encoded by the *LDHA* and *LDHB* genes, and are expressed in the skeletal muscle and heart. In addition, hetero-tetramers of the LDH-M and LDH-H subunits result in three, albeit less abundant, isoforms with distinct tissue distribution. Of note, LDHC is a testis-specific LDH that is composed of four LDH-C subunits. Substrate specificity of each LDH depends on its kinetic properties, with LDHA and LDHC preferentially converting pyruvate to lactate, whereas the reverse reaction is catalyzed by LDHB. LDHC is uniquely positioned as a potential anti-cancer target thanks to its restricted expression in normal somatic tissues and re-expression in many tumors (Koslowski *et al*, 2002). It has a well-established function in energy metabolism governing sperm motility and male fertility (Goldberg *et al*, 2010; Odet *et al*, 2008), however, its role in cancer is less understood.

Cui *et al* recently reported increased LDHC expression in tumor tissue and serum-derived exosomes of breast cancer patients, which correlated with poor survival, larger tumor size and recurrence (Cui *et al*, 2020). LDHC expression has also been positively correlated with shorter progression-free survival in renal cell carcinoma (Hua *et al*, 2017). Moreover, we recently demonstrated that LDHC is an immunogenic tumor associated antigen that can elicit a cytotoxic immune response against breast cancer cells (Thomas *et al*, 2020). One study demonstrated the presence of tumor-specific LDHC isoforms with defects in the structure of the catalytic domain that may result into non-functional, truncated splice variants (Koslowski *et al*., 2002). Others have reported that LDHC promotes cancer cell proliferation, migration and invasion in numerous cancer types (Chen *et al*, 2021; Hua *et al*., 2017; Kong *et al*, 2016). Given the role of a number of CTAs in maintaining genomic integrity during meiosis (McFarlane & Wakeman, 2017) and in particular of SSX2, NXF2, FMR1NB in cancer genomic instability (Gjerstorff *et al*., 2015), we investigated whether LDHC tumor re-expression might be related to the latter. We focused our study on basal-like breast cancer which is characterized by increased genomic instability (Cancer Genome Atlas, 2012; Manie *et al*, 2009; Shah *et al*, 2012). Using multiple breast cancer cell line models, we show that LDHC is involved in cell cycle regulation, and that targeting of LDHC improves treatment response to DNA damaging and DNA repair inhibiting agents.

## 2. RESULTS

### 2.1. LDHC expression in breast cancer

Analysis of the TCGA breast cancer dataset revealed a trend of increased *LDHC* expression in breast tumor tissue compared to normal tissue **(Figure 1A)**. The variability in expression levels prompted us to investigate *LDHC* expression across intrinsic molecular breast tumor subtypes. *LDHC* expression was significantly increased in basal-like tumors, the least favorable molecular subtype, compared to luminal A tumors, the most favorable subtype **(Figure 1B)**. Furthermore, high expression of *LDHC*, in particular in basal-like breast tumors, was associated with adverse overall (HR=1.33, p value=0.08) and disease-specific survival (HR=1.86, p value=0.002) **(Figure 1C)**. These findings are in accordance with a pro-tumorigenic role for LDHC in less favorable subtypes such as basal-like tumors.

**Figure 1.**
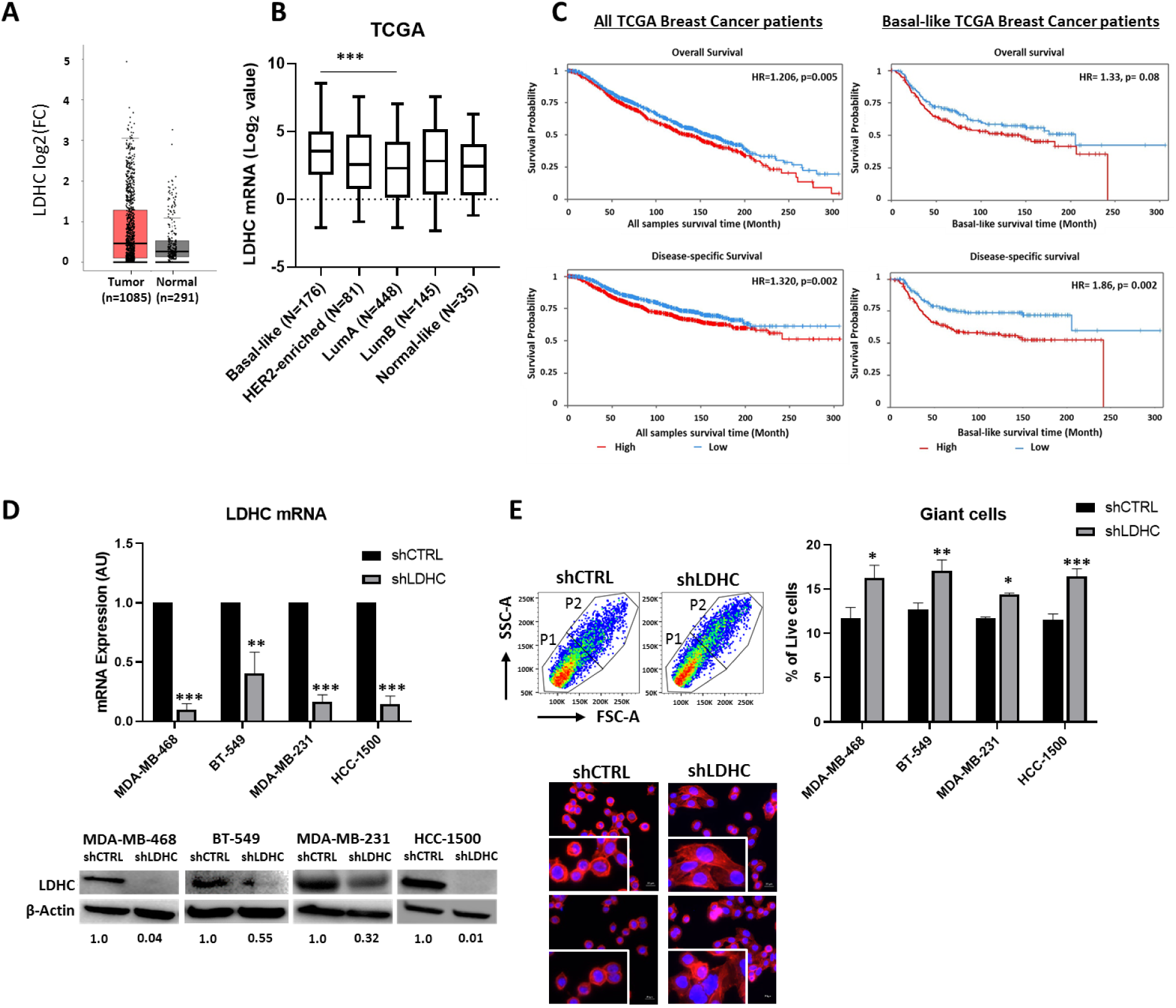
LDHC expression in breast cancer. A, Box plot of *LDHC* mRNA expression in tumor and normal breast tissue using the Breast Cancer (BRCA) TCGA dataset. B, Box plot depicting *LDHC* mRNA expression in intrinsic breast cancer subtypes in TCGA dataset. Statistical analysis performed using ANOVA and Tukey post hoc test, *** basal-like vs luminal A. C, Kaplan-Meier curves for overall and disease-specific survival, stratified by median *LDHC* mRNA expression. Hazards ratio (HR) and p values are indicated. D, *LDHC* mRNA, normalized to *RPLPO,* and LDHC protein expression in breast cancer cell lines stably transfected with shCTRL or shLDHC expression vectors. β-actin protein expression indicated as western blot loading control. Numbers under each lane represent mean densitometry values (arbitrary units) of LDHC signal normalized to β-actin from three independent experiments. E, (Top) Flow cytometry-based quantification of giant cells. Left-representative forward scatter (FSC-A) vs side scatter (SSC-A) flow cytometry plots for MDA-MB-468 cells with cell populations sub-grouped as P1 and P2 (giant cells). Right-frequency of giant cells (P2) in *LDHC*-silenced compared to control cells. (Bottom) representative immunofluorescence microscopy images of MDA-MB-468 with nuclear (DAPI, blue) and F-actin (red) staining (20x magnification), inserts at 2x zoom. Statistical analysis comparing shCTRL vs shLDHC performed using Student’s t-test. Error bars represent standard error of mean (±SEM) from three independent replicates. *p≤0.05, **p≤0.01, ***p≤0.001.

### 2.2. *LDHC* silencing induces the formation of giant cancer cells

In order to investigate the role of LDHC in breast cancer, its expression was stably silenced in three basal-like breast cancer cell lines (MDA-MB-468, BT-549, MDA-MB-231) alongside one non-basal, luminal A breast cancer cell line (HCC-1500). LDHC mRNA and protein expression was significantly reduced in all cell lines using LDHC-specific shRNA, albeit at different efficiencies, with the highest knockdown of *LDHC* in MDA-MB-468 and HCC-1500 and to a lesser extent in MDA-MB-231 and BT-549 **(Figure 1D).** Of note, no significant changes in LDHA and LDHB were observed upon *LDHC* silencing **(Supplementary Fig 1A)**. Silencing of *LDHC* significantly increased the number of giant cells and induced changes in the actin cytoskeleton whereby the ring-like distribution of filamentous actin (F-actin) stress fibers in shCTRL cells was replaced by a network of elongated actin filaments in a proportion of shLDHC cells **(Figure 1E)** (Vakifahmetoglu *et al*, 2008).

### 2.3. *LDHC* silencing induces genomic instability and mitotic catastrophe

Next, we investigated the presence and extent of genomic instability and nuclear aberrations as common features of giant cancer cells (Bakhoum & Compton, 2012; Cimini, 2008). We found that *LDHC* silencing increased the frequency and extent of polyploidy (≥4N) in MDA-MB-468 and BT-549 cells **(Figure 2A-B)** but not in breast cancer cells with inherent low levels of polyploidy (MDA-MB-231 and HCC-1500). MDA-MB-468 shLDHC cells displayed an increase in the proportion of cells with 8N, 10N and 12N polyploidy and decrease in 6N polyploidy **(Figure 2B)**. In addition, shLDHC cells displayed an increase in the presence of nuclear aberrations, including multinucleation (MNC), micronuclei (MN), nucleoplasmic bridges (NPB) and nuclear budding (NBUD) **(Figure 2C)**. As a countermeasure for increased genomic instability, cells can undergo mitotic catastrophe (MC), driving cells into an antiproliferative fate (Vitale *et al*, 2011). In line with this protective mechanism, a significant proportion of shLDHC cells exhibit mitotic catastrophe, which is associated with defects in spindle assembly **(Figure 2C)**. We were not able to validate these observations in the HCC-1500 cell line model due to the relatively small cell size impeding accurate microscopic assessment.

**Figure 2.**
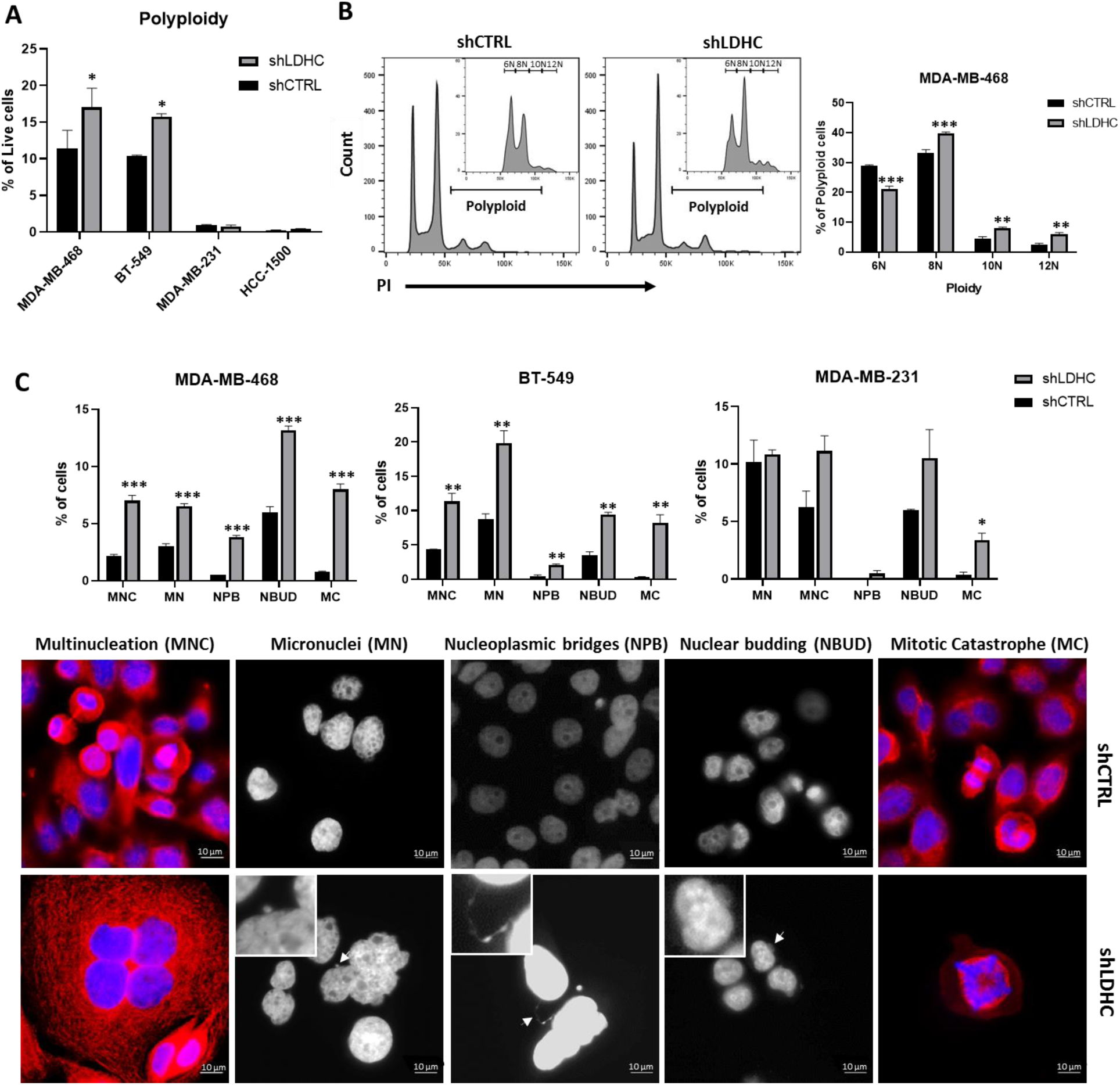
Silencing *LDHC* exacerbates polyploidy and nuclear aberrations. A, Frequency of cells exhibiting polyploidy (≥4N ploidy) in shLDHC versus shCTRL cells, as determined by PI flow cytometry analysis. B, Degree of polyploidy in shLDHC versus shCTRL MDA-MB-468 cells. Representative histograms with inserts representing genomic content of giant cells (P2). C, (Top) Frequency of nuclear aberrations in shLDHC and shCTRL cells (n=600). (Bottom) Representative immunofluorescence microscopy images of nuclear aberrations in MDA-MB-468 shCTRL and shLDHC cells with nuclear (DAPI, blue) and β-tubulin (red) staining (100x magnification), inserts at 2.5x zoom. All statistical analysis comparing shCTRL vs shLDHC performed using Student’s t-test. Error bars represent standard error of mean (±SEM) from three independent replicates. *p≤0.05, **p≤0.01, ***p≤0.001. MC-Mitotic catastrophe, MN-Micronuclei, MNC-Multinucleated cells, NBUD-Nuclear budding, NPB-Nucleoplasmic bridges.

### 2.4. *LDHC* silencing triggers excessive DNA damage and microtubule destabilization

In line with an increase in mitotic catastrophe, *LDHC* silencing markedly increased the expression of phospho-gamma-H2AX (γ-H2AX) in all four cell line models, indicating the presence of excess DNA damage **(Figure 3A)**. Assessment of homologous recombination (HR) and non-homologous end joining repair (NHEJ) pathway regulators demonstrated that both pathways were dysregulated by *LDHC* silencing, albeit to slightly different extents based on the cell line **(Supplementary Fig 1B).** Furthermore, *LDHC* silencing disrupted mitotic spindle organization, a common feature of cells undergoing mitotic catastrophe. More specifically, we found an increase in α-tubulin degradation **(Figure 3B)**, a decrease in its acetylation of lysine residue 40 **(Figure 3C)**, and a more punctate staining in shLDHC cells **(Figure 3E)**, collectively suggesting an increase in microtubule destabilization (Drubin *et al*, 1988; Janke & Montagnac, 2017; Kong *et al*, 1999). In addition, the expression of microtubule-associated protein 1B (MAP1B), involved in maintaining microtubule stability (Bhat & Setaluri, 2007), was significantly downregulated by *LDHC* silencing in three out of four cell lines **(Figure 3D-E)**. The lack of a decrease in *MAP1B* in MDA-MB-231 shLDHC cells suggests that alternative MAPs may be involved in driving microtubule destabilization in these cells. Together, these findings indicate that *LDHC* silencing promotes genomic instability and mitotic catastrophe concomitant with excessive DNA damage and mitotic spindle destabilization.

**Figure 3.**
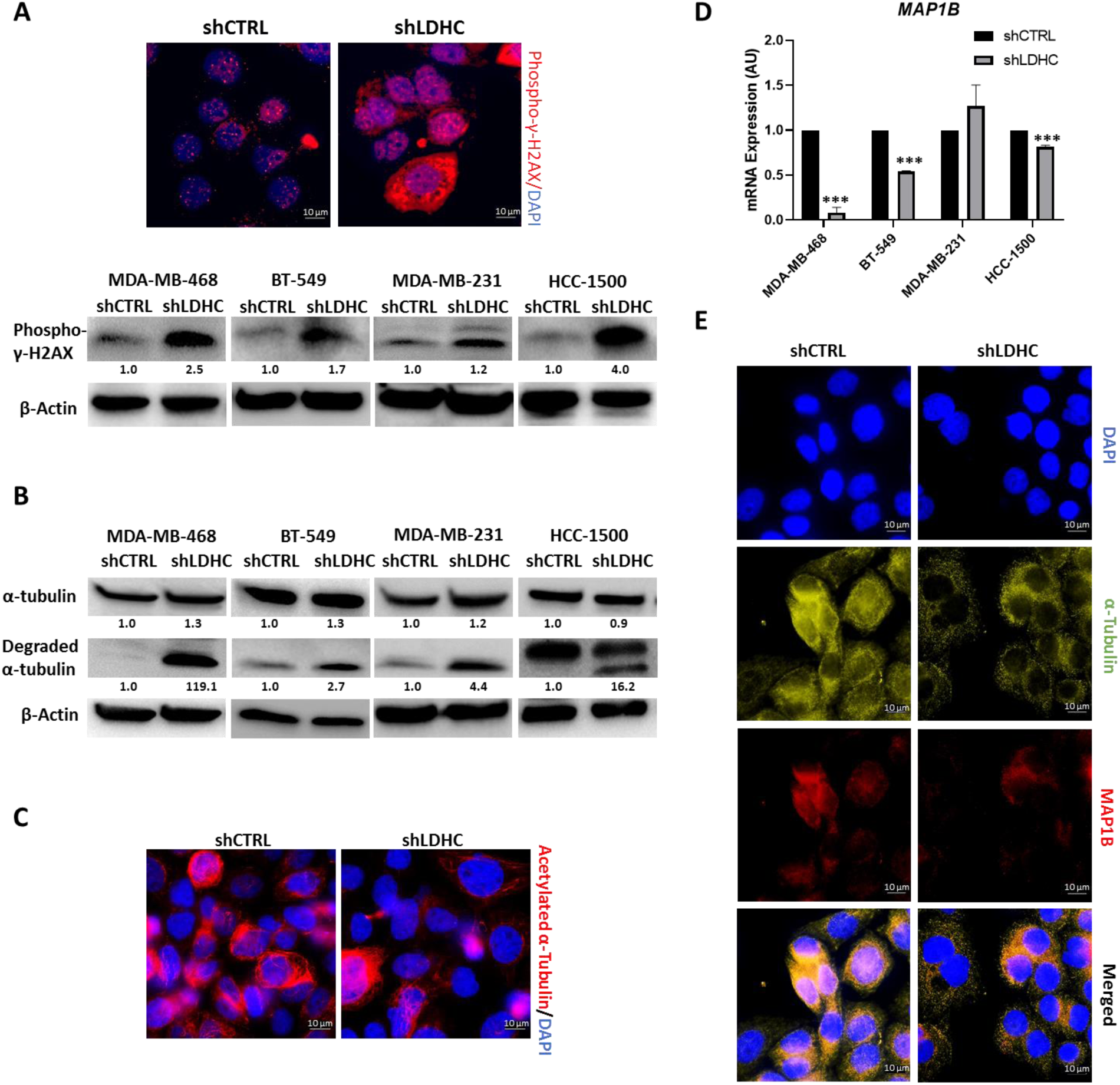
LDHC regulates DNA damage accumulation and microtubule network stability. A, (Top) Representative immunofluorescence microscopy images of phospho-γ-H2AX (red) and DAPI (blue)-stained nuclei in MDA-MB-468 cells (100x magnification). (Bottom) Western blot of phospho-γ-H2AX and β-actin as loading control. B, Western blot of full length and degraded α-tubulin expression. Note that HCC-1500 cells show an additional degraded product of α-tubulin with slightly higher molecular weight (not quantified). C, Representative immunofluorescence microscopy images of acetylated α-tubulin (red) and cell nuclei (DAPI, blue) in MDA-MB-468 cells (100x magnification). D, *MAP1B* mRNA expression, normalized to *RPLPO* expression. E, Representative immunofluorescence microscopy images of α-tubulin (green), MAP1B (red) and DAPI (blue)-stained nuclei in MDA-MB-468 cells (100x magnification). For panels A and B western blots, numbers under each lane represent mean densitometry values (arbitrary units) for respective protein signal normalized to β-actin from three independent experiments. Statistical analysis comparing shCTRL vs shLDHC performed using Student’s t-test. Error bars represent standard error of mean (±SEM) from three independent replicates. ***p≤0.001.

### 2.5. Mitotic catastrophe-associated cell fates

The activation of mitotic catastrophe can drive cells towards either of two cell fates; mitotic cell death or mitotic slippage. Hence, we assessed whether *LDHC* silencing facilitates either cell fate, and determined the expression of key players of mitotic entry and exit.

#### 2.5.1. Increased cell death

*LDHC* silencing induced apoptosis in all four breast cancer cell lines, as demonstrated by an increase in caspase 3 cleavage **(Figure 4A, Supplementary Fig 1C)**. Furthermore, flow cytometric analysis revealed an increase in the proportion of early (AnnexinV positive, PI negative) and/or late (AnnexinV positive, PI positive) apoptotic cells after *LDHC* silencing **(Figure 4A, Supplementary Fig 1D-E)**.

**Figure 4.**
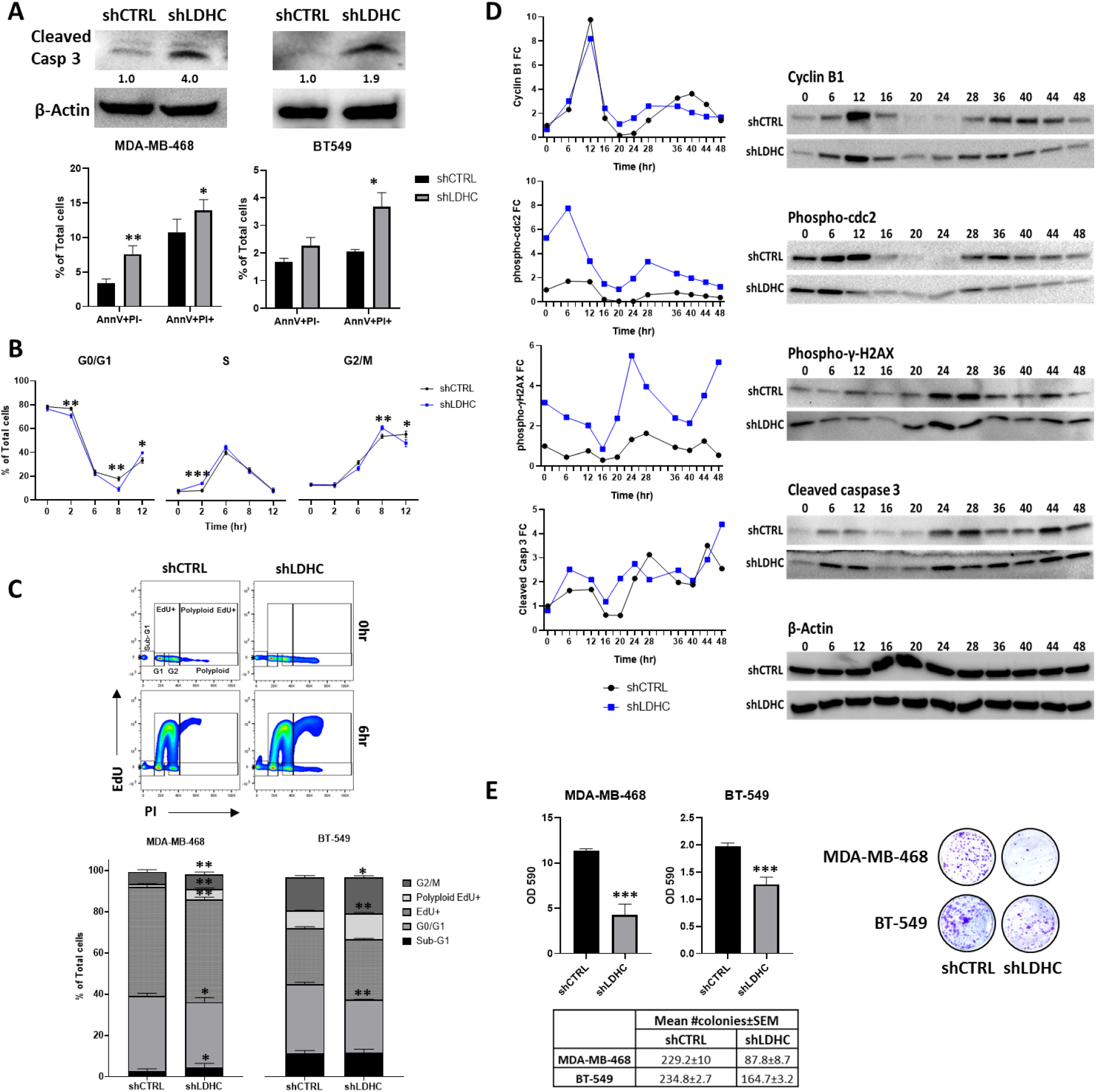
*LDHC* abrogation induces apoptosis, mitotic dysregulation and decreases long-term survival. A, (Top) Western blot of cleaved caspase 3 expression with β-actin as loading control. Numbers under each lane represent mean densitometry values (arbitrary units) for cleaved caspase 3 signal normalized to β-actin from three independent experiments. (Bottom) Quantification of apoptotic cells by AnnexinV/PI flow cytometry with early apoptosis defined as AnnV+PI- cells and late apoptosis as AnnV+PI+ cells. B, Time-course of cell cycle distribution of synchronized MDA-MB-468 cells using PI flow cytometry (error bars represent ± standard deviation). C, (Top) Representative cell cycle profile of asynchronous cell populations, as determined by EdU/PI flow cytometry. (Bottom) Proportion of asynchronous MDA-MB-468 and BT-549 cells in each cell cycle phase. D, Line chart representing the fold change (FC) in protein expression over time, as determined by western blot of cyclin B1, phopho-cdc2, phosph-γ-H2AX, cleaved caspase 3 expression with representative β-actin as loading control. Mean densitometry values (arbitrary units) of respective protein signals from three independent experiments were normalized to β-actin, and the normalized values were used to calculate the fold change in comparison to shCTRL at 0h. E, (Top left) Mean OD590 for crystal violet quantification, (Bottom left) mean number of colonies, and (Right) representative image as determined by clonogenic assay. All statistical analysis comparing shCTRL vs shLDHC performed using Student’s t-test. Error bars (in A, C, E) represent standard error of mean (±SEM) from three independent replicates. *p≤0.05, **p≤0.01, ***p≤0.001.

#### 2.5.2. Aberrant mitosis and loss of survival

Next, we investigated whether *LDHC* silencing is associated with mitotic slippage whereby cells prematurely exit mitosis by analyzing the cell cycle distribution of synchronized cells for up to 12 hours **(Figure 4B, Supplementary Fig 2A-B)**. In comparison to MDA-MB-468 shCTRL cells, shLDHC cells demonstrated rapid transition from G0/G1 to S and G2/M phase (2h and 8h respectively), and mitotic slippage with premature mitotic exit from the G2/M phase into the next cell cycle (12h) **(Figure 4B).** In addition, we found an increase in the proportion of shLDHC cells likely undergoing mitotic cell death (34% vs 21% sub-G1 shCTRL at 12h) **(Supplementary Fig 2C)**. Similarly, analysis of BT-549 shLDHC cells revealed rapid transition from G0/G1 to S phase **(Supplementary Fig 2B)** and increased cell death (12h) (**Supplementary Fig 2C**). Of note, mitotic slippage in BT-549 shLDHC cells was preceded by G0/G1 arrest in a likely attempt to repair DNA damage, and the cells displayed a significantly higher number of polyploid cells (12h), suggesting sustained defective mitosis **(Supplementary Fig 2B).** Both cell line models support the hypothesis that *LDHC* silencing induces mitotic dysregulation and subsequent apoptosis. Moreover, analysis of asynchronous populations of MDA-MB-468 and BT-549 cells **(Figure 4C)** provided further evidence for shLDHC-associated mitotic slippage with an increase in actively replicating polyploid cells (polyploid Edu+). Additional analyses of all four cell line models **(Figure 4C, Supplementary Fig 3A)** revealed a shLDHC-associated shift in the proportion of cells in G0/G1 (2N, EdU- negative) and G2/M (4N, EdU-negative) with MDA-MB-468 and HCC-1500 shLDHC cells also showing an increase in sub-G1 apoptotic cells. Together, these results suggest that silencing *LDHC* is associated with both mitotic slippage and arrest, followed by cell death.

To gain insight into the molecular mechanisms driving progression through mitosis, we assessed the activation status of the cyclin B1-cdc2 complex in the MDA-MB-468 cell line that demonstrated high *LDHC* silencing efficiency and a robust phenotype **(Figure 4D)**. Analysis of the expression of cyclin B1 and of inactive phosphorylated cdc2 (Tyr15 phospho-cdc2) revealed remarkable differences in expression dynamics in shLDHC cells compared to shCTRL cells over a time period of 48 hours (approximately two cycles of mitotic entry). Both control and *LDHC* silenced cells displayed the characteristic pre-mitotic peak in cyclin B1 expression at 12h post synchronization, however, shLDHC cells showed an earlier decline in phospho-cdc2 (6-16 vs 12-16h in shCTRL) in accordance with a more rapid G2/M transition and initiation of mitosis. Subsequently, shLDHC cells displayed an earlier second peak in cyclin B1 expression followed by cdc2 dephosphorylation (28 vs 40h in shCTRL), indicating that shLDHC cells prematurely entered the second round of mitosis. Of note, the earlier peak of cyclin B1 and phospho-cdc2 expression in shLDHC cells are followed by a slower rate of degradation. Moreover, their expression does not decline to baseline levels after the first round of mitosis (20-24 h). In concordance with our cell cycle profiling data, these observations suggest that *LDHC* silencing may result in two cell subpopulations, whereby one subset of cells undergoes mitotic slippage and the other experiences prolonged pre-mitotic arrest with likely subsequent cell death as a result of excessive, unrepaired DNA damage (Brito & Rieder, 2006; Zeng *et al*, 2019). Indeed, it is well known that the balance of two opposing networks, involving cyclin B1 degradation and caspase cleavage, determine mitotic cell fate (Topham & Taylor, 2013). Analysis of phospho-γ-H2AX levels demonstrated a high level of DNA damage in shLDHC cells at 0 h and at the mitotic phases (24 h and 48 h) **(Figure 4D)**. Furthermore, assessment of caspase 3 cleavage **(Figure 4D)** provides evidence of a sustained increase in shLDHC cell death throughout the cell cycle (6-12h and 20h onwards) and in subsequent cell cycles (48h).

In addition to undergoing apoptosis, mitotic catastrophe can also drive cells to enter a state of senescence as observed in HCC-1500 shLDHC cells but not in the other three cell lines **(Supplementary Fig 3B).** Strikingly, silencing of *LDHC* significantly decreased the colony-formation ability, hence long-term survival, of all four breast cancer cell lines **(Figure 4E, Supplementary Fig 3C)**.

### 2.6. *LDHC* silencing dysregulates multiple cell cycle checkpoints

Based on our observations of increased mitotic dysregulation by *LDHC* silencing, we explored potential defects in G1/S, intra-S, G2/M and Spindle Assembly checkpoint (SAC) regulators using the MDA-MB-468 cell line model. Differential gene expression and gene ontology analysis of 84 cell cycle-related genes revealed an enrichment of genes involved in dysregulated DNA damage response and impaired mitotic fidelity in *LDHC* silenced cells **(Figure 5A, Supplementary Table 2)**.

**Figure 5.**
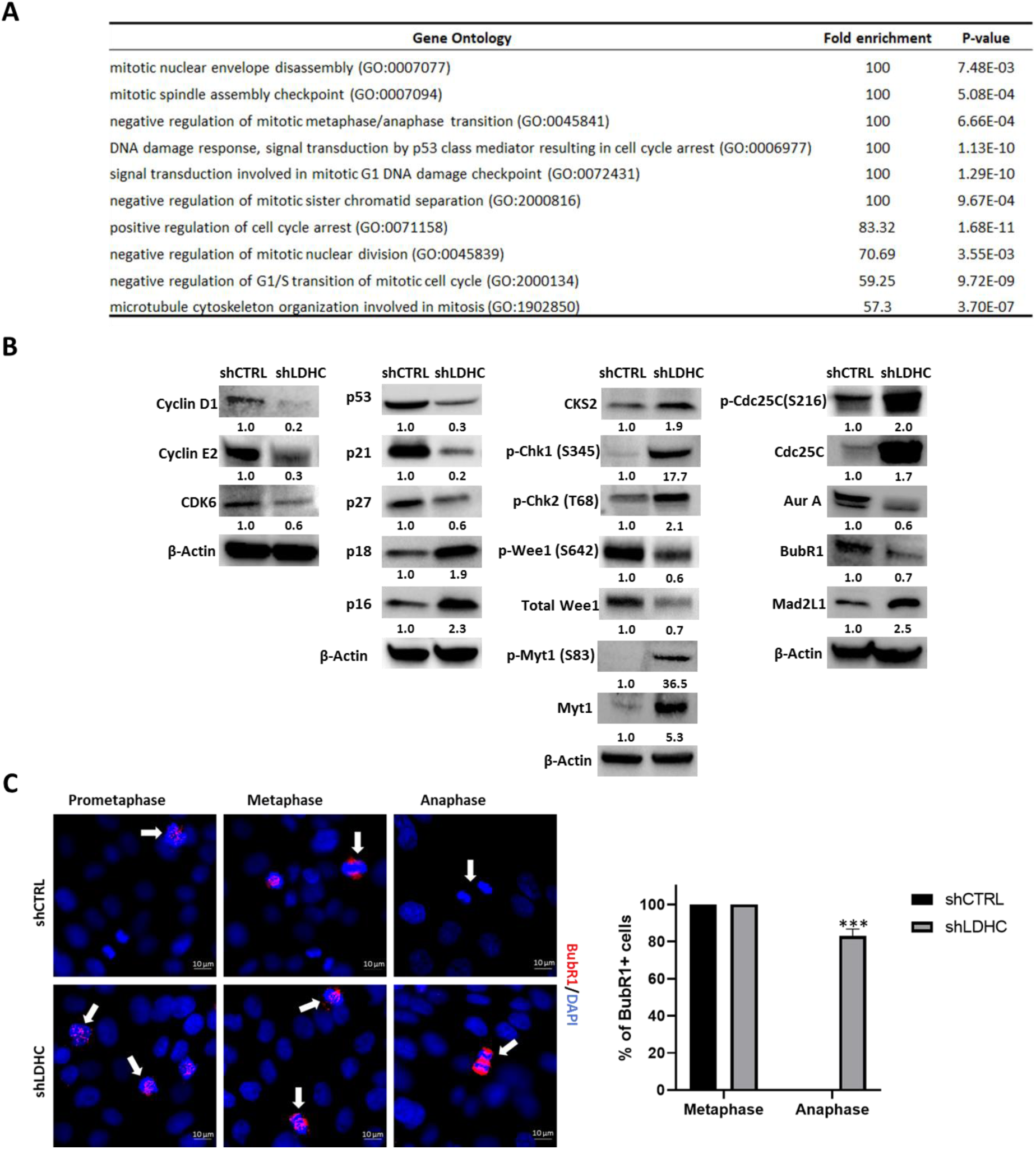
LDHC and cell cycle regulation of MDA-MB-468 cells. A, GO enrichment of differentially expressed genes after *LDHC* silencing as determined by the Cell Cycle RT2 Profiler qPCR array. B, Western blot of cell cycle regulator protein expression with β-actin as loading control. Numbers under each lane represent mean densitometry values (arbitrary units) for respective protein signal normalized to β-actin from three independent experiments. C, Representative immunofluorescence microscopy images of BubR1 (red) and DAPI-stained nuclei (blue) in prometaphase, metaphase and anaphase stages of mitosis (indicated by white arrows, 100x magnification). All statistical analysis comparing shCTRL vs shLDHC performed using Student’s t-test. Error bars represent standard error of mean (±SEM) from three independent replicates with n=90 cells for each condition. ***p≤0.001.

Furthermore, we found that *LDHC* silencing altered the protein expression of cell cycle regulators at multiple checkpoints. For instance, the expression of the G1/S checkpoint regulators Cyclin D1, Cyclin E2 and Cyclin-dependent kinase (CDK)-6 was downregulated in shLDHC cells **(Figure 5B),** suggesting that less cells reside in the G1 phase or that the G1 phase is shortened, which is in line with our cell cycle profile observations **(Figure 4C)**. The observed upregulation of INK4 type CDK inhibitors (p16, p18) and associated downregulation of p53 and its downstream targets p27 and p21 (Mirzayans *et al*, 2012) further supports the likelihood of G1/S checkpoint dysregulation in shLDHC cells. In addition, *LDHC* silencing increased the expression of cyclin-dependent kinase subunit 2 (CKS2), likely mediating an override of the intra-S phase checkpoint in the presence of replication stress (Liberal *et al*, 2012) and facilitating transition from G2 to M phase (Martinsson-Ahlzen *et al*, 2008) **(Figure 5B, Supplementary Table 2)**. Expression analysis of the negative G2/M checkpoint regulators Wee1 and Myt1 demonstrated a decrease in total and active phosphorylated Wee1 (Ser642), and conversely an increase in total and inactive phosphorylated Myt1 (Ser83) in shLDHC cells, supporting our previous observations that *LDHC* silencing promotes mitotic entry. On the other hand, we found an increase in phosphorylated checkpoint proteins Chk1 (Ser345) and Chk2 (Thr68), indicating the presence of single- and double-strand DNA breaks, that in turn inactivate Cdc25C by phosphorylation at Ser216, resulting in G2/M arrest. Collectively, these findings corroborate the presence of two shLDHC cell subpopulations; one population that undergoes G2/M arrest and another that undergoes checkpoint adaptation and slippage (Swift & Golsteyn, 2016). Further evidence for the existence of two shLDHC cell populations was provided by the assessment of mitotic/SAC regulators such as Aurora A (Roghi *et al*, 1998; Seki *et al*, 2008) and Mad2L1. We found that total and phosphorylated Aurora A expression were significantly reduced in shLDHC cells **(Figure 5B, Supplementary Table 2)**, and as such may result in the loss of an active SAC with premature transition from metaphase into anaphase (Courtheoux *et al*, 2018). In addition, expression of Mad2L1 was upregulated after *LDHC* silencing, possibly triggering cells to undergo long-term SAC activation followed by mitotic slippage (Raab *et al*, 2015) **(Figure 5B, Supplementary Table 2)**. In contrast, we observed a slight decrease in the expression of the kinetochore-associated protein BubR1 in shLDHC cells, thus impairing accurate spindle attachment and chromosome segregation. Although the cellular localization of BubR1 did not differ in the pro-metaphase between shCTRL and shLDHC cells, its expression at the kinetochores and at segregating chromosomes in the metaphase and early anaphase was dysregulated in shLDHC cells, indicating defective sister chromatid segregation and prevention of anaphase onset **(Figure 5C)**. In line with this, the majority of shLDHC cells in anaphase remained BubR1 positive whereas shCTRL cells completely lost BubR1 expression going from metaphase to anaphase.

In conclusion, assessment of the expression of cell cycle regulators in the MDA-MB-468 cell line confirmed the presence of two cell subpopulations after *LDHC* silencing: one subpopulation undergoing mitotic arrest at the G2/M and/or SAC checkpoint, and another undergoing mitotic slippage. This phenotype of mitotic dysregulation upon *LDHC* silencing was observed in all four breast cancer cell lines, however, the molecular mechanistic underpinnings of these observations may vary between cell lines based on their genetic landscape.

### 2.7. *LDHC* silencing sensitizes cancer cells to DNA damage repair inhibitors and DNA damage inducers

DNA damage repair inhibitors and DNA damage inducing agents, including the Poly ADP-ribose polymerase (PARP)-inhibitor olaparib and cisplatin, are widely used anti-cancer drugs with particular benefit to patients with defects in DNA damage response pathways such as BRCA1/2-positive or ‘BRCAness’ basal breast cancer patients (Byrum *et al*, 2019; De Summa *et al*, 2013). Since we demonstrated that *LDHC* silencing increases DNA damage accumulation in breast cancer cells with alterations in the expression of both HR and NHEJ regulators, we explored the sensitivity of shLDHC cells to olaparib and cisplatin. Treatment with either drug further induced apoptosis in giant shLDHC cells compared to shCTRL cells **(Figure 6A, Supplementary Fig 4A-C)**. In accordance, olaparib and cisplatin treatment augmented the expression of cleaved caspase 3 in shLDHC cells **(Figure 6B)**. Moreover, we observed a significant increase in DNA damage after olaparib and cisplatin treatment, albeit at similar levels in both shCTRL and shLDHC treated cells **(Figure 6C)**. However, shLDHC cells with excessive DNA damage displayed apoptotic nuclear features such as highly condensed DNA and nuclear fragmentation. Finally, the colony formation ability of shLDHC cells was further compromised after treatment with olaparib and cisplatin compared to shCTRL treated cells **(Figure 6D, Supplementary Fig 4D)**.

**Figure 6.**
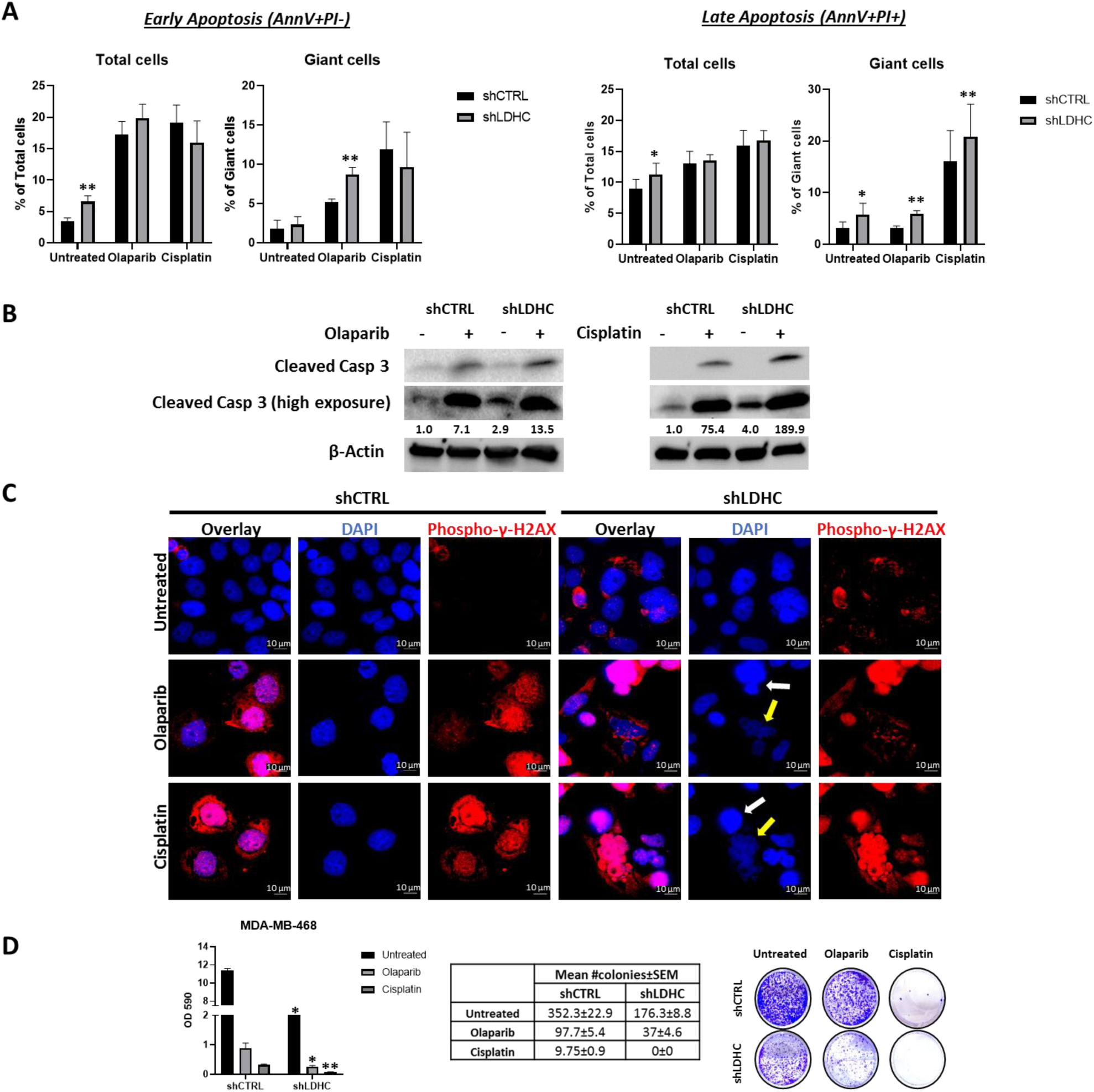
*LDHC* silencing improves sensitivity of MDA-MB-468 cells to DNA damage inducers and DNA damage repair inhibitors. A, Annexin V/PI flow cytometric quantification of apoptosis after 72hrs of treatment with olaparib or cisplatin. B, Western blot of cleaved caspase 3 protein expression with β-actin as loading control. Numbers under each lane represent mean densitometry values (arbitrary units) for cleaved caspase 3 signal (high exposure blot) from three independent experiments normalized to β-actin. C, Representative immunofluorescence microscopy images of phospho-γ-H2AX (red) and DAPI-stained nuclei (100x magnification). Yellow arrows indicate early apoptotic cells and white arrows indicate late apoptotic cells. D, (Left) Mean OD590 for crystal violet quantification, (Middle) mean number of colonies and (Right) representative images taken at 14 days of culture post-treatment (72h). All statistical analysis comparing shCTRL vs shLDHC performed using Student’s t-test. Error bars represent standard error of mean (±SEM) from three independent replicates. *p≤0.05, **p≤0.01

## 3. DISCUSSION

Cancer testis antigens, characterized by tumor-restricted expression and immunogenicity, are attractive candidate targets for cancer therapy. Gaining insight into their role in tumorigenesis could facilitate the identification of individual highly tumor-specific CTAs with therapeutic potential that may synergistically improve the efficacy of available cancer therapies (Gibbs & Whitehurst, 2018). While cancer cells commonly exhibit deficiencies in DNA damage repair pathways, there appears to be a threshold whereby low levels of genomic instability drive tumorigenicity while excess genomic aberrations compromise cellular fitness through cell cycle arrest, senescence or cell death (Bakhoum & Compton, 2012). Here, we report that targeting of LDHC in breast cancer cells has the potential to tip the fine balance between tolerable and excessive levels of genomic damage in favor of the latter, and as such to sensitize cancer cells to DNA damage inducers and DNA damage repair inhibitors **(Figure 7)**.

**Figure 7.**
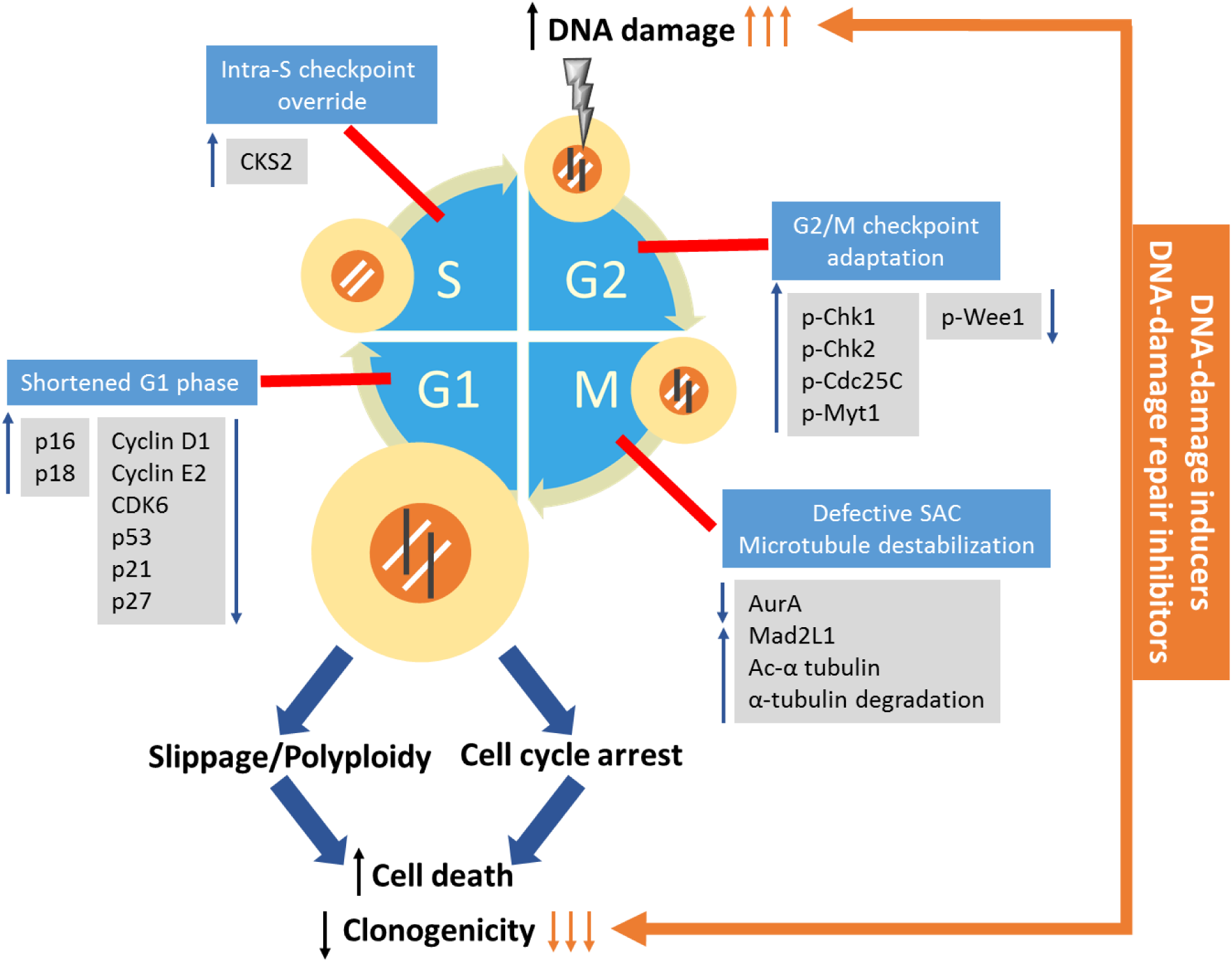
Schematic model of the effect of LDHC targeting on cell cycle checkpoints and cell fate. Silencing of *LDHC* in breast cancer cells increases the rate of mitotic entry by dysregulation of several cell cycle checkpoints, resulting in a shorter cell cycle. Firstly, targeting *LDHC* shortens the G1 phase and downregulates several molecules controlling the G1/S checkpoint and mediates override of the intra S-phase checkpoint. Next, G2/M checkpoint regulators are aberrantly expressed in *LDHC* silenced cells. Together, this results in excessive DNA damage (black lines) due to the lack of functional repair mechanisms. On mitotic entry, the unstable microtubule network in *LDHC* silenced cells triggers defective chromosome segregation, thus activating the spindle assembly checkpoint (SAC). While a proportion of cells with high levels of genomic instability undergo arrest and cell death, an additional population of cells with lower levels of instability undergo long-term SAC activation and mitotic slippage in the absence of cytokinesis. These cells form giant polyploid cancer cells that subsequently undergo cell death or senescence, ultimately diminishing clonogenicity or long-term survival of cancer cells. Additionally, targeting *LDHC* in combination with DNA-damaging/DNA-repair inhibiting agents (orange arrows) synergistically dysregulates the DNA damage repair pathways and promotes cell death pathways, resulting in significant loss of cell survival. The G1/S, intra-S, G2/M and SAC checkpoints are depicted as red lines. The molecular regulators depicted were found to be involved in *LDHC* silenced MDA-MB-468, however, we cannot exclude that there may be cell-line dependent differences in regulators.

Through a comprehensive analysis of four breast cancer cell lines, we demonstrated that *LDHC* silencing induces DNA damage accumulation and impairs mitotic fidelity by dysregulating multiple cell cycle checkpoints and microtubule assembly, ultimately resulting in mitotic catastrophe, apoptosis and reduced long-term survival **(Figure 7, Supplementary Table 3)**. Of note, these observations were consistent in three basal-like breast cancer cell lines and in one luminal A cell line, indicating the broad applicability of LDHC function in maintaining mitotic fidelity. Although the various cell lines share similar cell fates upon *LDHC* silencing, their genetic background varies and the specific molecular mediators and mechanisms involved may be cell line-dependent. In conjunction with *LDHC* knockdown efficiency, differences in genetic mutational status may partly explain some of the mechanistic differences observed in the various cell lines. For instance, the genetic landscapes of the cell lines differ in the mutational status of the Rb and p53 tumor suppressor genes that regulate the G1/S checkpoint **(Supplementary Table 3)**. As such, cell lines with loss of Rb and/or mutant p53 rely more heavily on a functional G2/M DNA damage checkpoint for tumor propagation, which may be reflected in molecular dysregulation nuances induced by *LDHC* silencing (Blandino & Di Agostino, 2018). Furthermore, we show in p53 mutant MDA-MB-468 cells that *LDHC* silencing reduced the expression of the oncogenic, mutant form of p53, thus contributing to the loss-of-survival phenotype (Lim *et al*, 2009). Conversely, HCC-1500 cells lack p53 expression and display a more moderate phenotype upon *LDHC* silencing. This raises the question whether breast tumors with oncogenic p53 mutations that express high levels of *LDHC* **(Supplementary Fig 5)** could be more susceptible to LDHC targeting through disruption of the G2/M and SAC checkpoints.

Interestingly, although none of the four cell lines in our study harbor *BRCA* mutations, we observed DNA damage accumulation upon *LDHC* silencing suggesting that DNA repair pathways are likely impaired. A more in-depth analysis of key molecules involved in DNA repair signaling revealed dysregulation of both the homologous recombination and non-homologous end joining repair pathways in *LDHC* silenced cells. In addition, *LDHC* silencing in MDA-MB-468 cells, which display a greater *LDHC* knockdown efficiency and a robust phenotype, downregulated the expression of Aurora A and its downstream mediator Wee1 and inactivated Myt1. As such, *LDHC* silencing induced disruption of the G2/M checkpoint and homologous recombination repair pathway, allowing cells to proceed into mitosis with excess unrepaired DNA damage (Karnak *et al*, 2014; Krajewska *et al*, 2013). Moreover, Aurora A plays a critical role in microtubule nucleation and elongation indicating that its reduced expression, together with the loss of the microtubule-associated protein MAP1B, is potentially involved in the *LDHC* silencing-induced microtubule instability (Willems *et al*, 2018). In addition to DNA damage accumulation attributed to dysregulated mitosis and DNA repair pathways, it remains to be determined if *LDHC* silencing affects the levels of DNA damage inducers such as reactive oxygen species or impairs the functionality of NAD+-dependent enzymes such as PARPs and sirtuins, involved in maintaining genomic stability, through modulation of pyruvate-lactate interconversion and consequently NAD+ and NADH levels (Bosch-Presegue & Vaquero, 2014; Gupte *et al*, 2017).

Given the increase in DNA damage and dysregulation of DNA damage repair pathways upon *LDHC* silencing, we speculated that, in analogy with treatment response of BRCA-deficient tumor cells, *LDHC* silencing could improve the response to DNA damage repair inhibitors and/or DNA damage inducers that rely on DNA damage-induced mitotic catastrophe to trigger cell death (Farmer *et al*, 2005; Mc Gee, 2015). Indeed, we found that silencing *LDHC* sensitizes breast cancer cells to olaparib and/or cisplatin, thereby improving the efficacy of either treatment with subtle differences in cell line sensitivity.

Our findings indicate that targeting of LDHC likely interferes with tumor cell survival by a multitude of mechanisms affecting genomic stability, including dysregulated cell cycle progression. To date, several cell cycle inhibitors are currently under investigation or are implemented in breast cancer management, including inhibitors against Wee1 (Pitts *et al*, 2020), Aurora A (Melichar *et al*, 2015), CDK6 (Alvarez-Fernandez & Malumbres, 2020), mutant p53 (Mantovani *et al*, 2019) as well as microtubule inhibitors (Jordan & Wilson, 2004). However, treatment with these inhibitors as single agents shows low response rates and results in the development of acquired resistance (Alvarez-Fernandez & Malumbres, 2020; Jordan & Wilson, 2004; Lewis *et al*, 2019). Of note, we found that targeting of *LDHC* negatively impacts each of these cell cycle regulators **(Figure 7)**, indicating that LDHC may prove to be a therapeutically superior target over individual cell cycle molecule inhibitors. Moreover, the tumor-specific expression of LDHC allows precise targeting of cancer cells with limited off-target effects, which is an improvement over currently available cell cycle inhibitors that affect any dividing somatic cells. Further, the combination effect of LDHC abrogation and DNA damaging agents/PARPi may allow reducing the drug dosage of the latter thereby lowering the risk of toxic side effects such as myelosuppression and nephrotoxicity. Although promising, the observations from this study remain to be confirmed in *in vivo* models to corroborate the notion of targeting LDHC to improve clinical outcome of breast cancer patients who may benefit from treatment with DNA damaging agents or PARPi, currently only prescribed to *BRCA* mutant patients who constitute a mere 9-15% of TNBCs (Armstrong *et al*, 2019). Notably, non-commercial oxamic acid analogues with high affinity and potency against LDHC have been described, however these have not yet been investigated for their anti-cancer properties (Rodriguez-Paez *et al*, 2011).

To conclude, we believe that developing LDHC-specific therapeutic interventions using specific chemical inhibitors or siRNA delivery approaches presents an attractive paradigm to enhance the efficacy of current drugs and to improve clinical outcome in breast cancer.

## 4. MATERIALS & METHODS

### 4.1. Transcriptomic and survival analysis

The Gene Expression Profiling Interactive Analysis (GEPIA) platform (http://gepia.cancer-pku.cn/index.html) was used to visualize *LDHC* transcriptomic data in normal and breast cancer tissue from The Cancer Genome Atlas (TCGA) and the Genotype-Tissue Expression (GTEx) datasets. The expression of *LDHC* was calculated in transcripts per million mapped reads (TPM), log-transformed for differential analysis and represented by the log2 fold change (log2FC) between tumor and matched normal data (median value of tumor samples minus median value of normal samples). The Breast Cancer Integrated Platform (BCIP) (http://www.omicsnet.org/bcancer/database) (Wu *et al*, 2017) was utilized to visualize *LDHC* expression in the TCGA intrinsic breast cancer subtypes and to generate Kaplan–Meier survival curves stratified by *LDHC* high versus low expression (cutoff=median log2 value).

### 4.2. Cell culture

All breast cancer cell lines were acquired from the American Tissue Culture Collection (ATCC) and authenticated by Short Tandem Repeat (STR) analysis. MDA-MB-468 and MDA-MB-231 cells were cultured in Dulbecco’s Minimum Essential Media (Gibco) supplemented with 10% v/v fetal bovine serum (FBS) (Hyclone US origin, GE Life Sciences), 50 U/mL Penicillin and 50 µg/mL Streptomycin (Gibco). BT-549 cells were maintained in ATCC-formulated Roswell Park Memorial Institute (RPMI) 1640 medium (Gibco) supplemented with 10% (v/v) FBS (Hyclone US origin, GE Lifescience), 50 U/mL penicillin and 50 µg/mL streptomycin (Gibco), and 0.023 IU/mL insulin (Sigma-Aldrich). HCC-1500 cells were cultured in RPMI 1640 media (Gibco) supplemented with 10% FBS, 50 U/mL Penicillin and 50 µg/mL Streptomycin (Gibco). All cell lines were maintained in a humidified incubator at 37°C and 5% CO2. Regular mycoplasma testing was conducted using a polymerase chain reaction (PCR)-based detection assay. Early passage cells (<P10) were used for all experiments.

### 4.3. Stable silencing of LDHC

All cell lines were transduced at 70-80% confluency with purified LDHC-specific shRNA lentiviral particles (SMARTvector Lentiviral Human LDHC hCMV-TurboGFP shRNA, Dharmacon, V3SH11240-227292996/229943916) or scrambled negative control lentiviral particles (SMARTvector Non-targeting hCMV-TurboGFP Control Particles, Dharmacon, #S-005000-01) and polybrene transfection reagent (Millipore). The packaged viral vectors encode green fluorescent protein (GFP) reporters and a gene conferring resistance to Puromycin antibiotic. Six days post-transduction, transduced cells were selected in complete growth media, supplemented with 0.5 µg/mL Puromycin (Sigma-Aldrich).

### 4.4. Quantitative real-time PCR

Total RNA was extracted using Tri reagent (Ambion) as described previously (Naik *et al*, 2017). Reverse transcription of RNA was performed using MMLV-Superscript (Thermo Fisher Scientific) and random hexamers. *LDHC* expression was quantified by specific 5′FAM-3′MGB Taqman gene expression primer/probe sets (Hs00255650_m1, Applied Biosystems). *MAP1B* expression was quantified using primers for SYBR-based qPCR (F: CACCTCGCCTAGCCTGTC, R: CGGATTCCGAGCTCGATG), designed using PrimerBLAST (NCBI), and PowerUp SYBR Green mastermix (Applied Biosystems). qRT-PCR was performed on the QuantStudio 7 system (Applied Biosystems). Relative expression levels were normalized to the housekeeping gene *RPLPO* (Taqman primer/probe 4333761F or SYBR primers F: TCCTCGTGGAAGTGACATCG, R: TGGATGATCTTAAGGAAGTAGTTGG).

### 4.5. Immunoblotting

Cell protein lysate was isolated using RIPA buffer (Pierce) supplemented with HALT protease and phosphatase inhibitor cocktail (Thermo Fisher Scientific). Western blotting was performed using a standard protocol as previously described (Al-Khadairi *et al*, 2019). Primary antibodies utilized are listed in **Supplementary Table 1**. Horseradish Peroxidase (HRP)-linked anti-rabbit/mouse secondary antibody incubation followed by enhanced chemiluminescent substrate (ECL) Supersignal West Femto (Pierce) incubation was used to visualize the protein bands of interest on the ChemiDoc XRS+ Imaging system (Biorad). Images acquisition and densitometry analysis was performed using the Image Lab software (Biorad).

### 4.6. Immunofluorescence Microscopy

Immunofluorescence staining was performed according to standard protocols as described previously (Al-Khadairi *et al*., 2019). Briefly, 80,000 cells were plated on Poly-Lysine coated glass coverslips (Corning), fixed with 4% paraformaldehyde (ChemCruz) and permeabilized with 0.1% Triton X-100 (Sigma). Primary antibodies against β-tubulin (Cell Signaling), phospho-γH2AX (Abcam), alpha-tubulin (Li-Cor), acetylated alpha-tubulin (Cell Signaling), MAP1B (Abcam), BubR1 (Abcam), and Alexa Fluor 568 phalloidin (Thermo Fisher Scientific) are listed in **Supplementary Table 1**. Fluorescently-labelled secondary antibody anti-rabbit/mouse Alexa Fluor 555/647 (Thermo Fisher Scientific) were used. Cell nuclei were counterstained with 4′,6-diamidino-2-phenylindole (DAPI) (Thermo Fisher Scientific) and cells were mounted using Prolong Gold Antifade reagent (Invitrogen). Images were captured using an upright fluorescent microscope (Zeiss Axioimager, 40x or 100x oil objective) and the Zen Pro 2 acquisition and analyses software. The frequency of nuclear aberrations in cells was determined by manual quantification (100x magnification) of 600 DAPI-stained nuclei per condition (approximately 120 exclusive fields), and BubR1-staining was assessed in 90 DAPI-stained nuclei for each condition (approximately 15 exclusive fields).

### 4.7. Flow cytometry

Cells were harvested by centrifugation and washed in cold phosphate buffered saline (PBS). For cell cycle analysis and enumeration of polyploid cells, cells were fixed with ice-cold 66% ethanol overnight at 4°C and stained with propidium iodide (PI)/RNase A (Abcam) for 30 min at 37°C in the dark. For AnnexinV/PI quantification, cell were resuspended in 1x Annexin binding buffer (Thermo Fisher Scientific) and stained with AnnexinV BV421 (BD Biosciences) and PI at room temperature for 15 min. Flow cytometry analyses were performed using the LSRFortessa X-20 system and FlowJo software (BD Biosciences).

### 4.8. EdU incorporation assay

Cells were seeded in six-well plates and incubated for 6 h with 10 µM of the thymidine analog 5-Ethynyl-2′-deoxyuridine (EdU) in complete growth media. Next, cells were centrifuged, washed twice with PBS and fixed with ice-cold 66% ethanol overnight at 4°C. The fixed cells were brought to room temperature, washed twice with PBS and stained with the Click-IT® EdU Alexa Fluor® 647 Flow Cytometry kit as per manufacturer’s instructions (Applied Biosystems). Cells were finally stained with PI and analyzed by flow cytometry.

### 4.9. Cell cycle synchronization

Cells were seeded in a six-well plate and synchronized at G1/S phase by a double thymidine block (Chen & Deng, 2018). Briefly, cells were first treated with 2 mM thymidine (Sigma-Aldrich) for 18 h, followed by two washes with PBS and a 9 h release in normal growth media. The cells were then treated with a second thymidine block (2mM) for 16 h. Subsequently, cells were washed twice with PBS and incubated in complete growth media. Cells were harvested at multiple timepoints for cell cycle analysis (PI flow cytometry) and western blot analysis.

### 4.10. Clonogenic assay

A total of 10^3^ cells/well were seeded in six-well plates and maintained in complete growth media for 14 days after which the cells were washed with PBS and stained with crystal violet (5% Bromophenol blue, 25% methanol) for 20 min at 37°C. Excess stain was washed away with distilled water. Stained colonies were first counted manually and then the stain within the colonies was solubilized using 10% sodium dodecyl sulfate (SDS), followed by measurement of the optical density at wavelength 590 nm.

### 4.11. Senescence assay

A total of 5×10^5^ cells/well were seeded in six-well plates and stained for senescence-associated ß-galactosidase (SA-ß-Gal) activity using the cellular senescence assay kit (Cell Biolabs, Inc.) according to manufacturer’s instructions. Cells were incubated with staining solution overnight (MDA-MB-468, BT-549, MDA-MB-231) or for 4 h (HCC-1500) and visualized using an inverted light microscope (Zeiss Primo Vert, 40x objective) and the Zen Pro 2 acquisition and analysis software.

### 4.12. EMT RT2 Profiler™ PCR array

Differential expression of cell cycle-associated genes was determined using the EMT RT2 profiler qPCR array (Qiagen, PAHS-020Z) and analyzed using the Qiagen online Data Analysis Tool. Expression was normalized to the housekeeping gene *RPLPO*, and the threshold for differential expression was set at absolute log2 fold change ≥ 2.

### 4.13. Gene Ontology enrichment analysis

Enrichment analysis of gene ontology (GO) was performed using the PANTHER (Protein Analysis Through Evolutionary Relationships) online tool (http://www.pantherdb.org) (Mi *et al*, 2019). The Fisher’s Exact test with Bonferroni correction was used to identify the enriched GO biological process with p ≤ 0.05 and fold-enrichment > 50%.

### 4.14. Treatment with DNA-damaging agents

Cells were seeded in six-well plates and treated with cisplatin (Sigma Aldrich) or olaparib (Selleck Chemicals). The drug IC50 dosages for each cell line were identified from the Genomics in Drug Sensitivity in Cancer online database (https://www.cancerrxgene.org/) (Yang *et al*, 2013) and verified in-house using the WST-1 assay (Abcam) after 72 h treatment with the drug. The IC50 drug dosages used for the various cell lines were as following: 30 µM olaparib and 4 µM cisplatin for MDA-MB-468, 210 µM olaparib and 10 µM cisplatin for BT-549, 28.8 µM olaparib and 19 µM cisplatin for MDA-MB-231, and 111 µM olaparib and 140 µM cisplatin for HCC-1500. Cells were treated for 72 h in complete growth media followed by western blot analyses, clonogenic assay, AnnexinV/PI flow cytometry or immunofluorescence microscopy.

### 4.15. Statistical analysis

Normality of data was assessed using the Shapiro–Wilk test. Non-parametric analyses were conducted using Kruskal–Wallis test, while parametric analyses were performed using Student’s t-test or analysis of variance (ANOVA). P value ≤0.05 was defined as statistically significant. Data are represented as mean ± standard error of mean (SEM) of at least three independent biological replicates. Statistical analyses and data representation were performed using GraphPad Prism v8.0.0.

## ACKNOWLEDGEMENTS

This work was supported by a grant from the Qatar Biomedical Research Institute (Grant ID #2016-003, VR94), Qatar Foundation awarded to Dr Julie Decock. We would like to thank Dr Hirohito Yamaguchi for providing olaparib and Dr Wouter Hendrickx for providing cisplatin. We are grateful to Dr Remy Thomas for her laboratory assistance.

## AUTHOR CONTRIBUTIONS

**Adviti Naik**: Conceptualization, Methodology, Validation, Formal analysis, Writing - Original Draft, Visualization; **Julie Decock**: Conceptualization, Writing - Review & Editing, Supervision, Project administration, Funding acquisition.

## DECLARATION OF INTEREST

The authors declare no conflicts of interest.

## Abbreviations

ANOVA: Analysis of variance
ATM: Ataxia-Telangiectasia Mutated
ATR: Ataxia telangiectasia and Rad3 related
CDK: Cyclin Dependent Kinase
CTA: Cancer Testis Antigen
DNA-PKcs: DNA-dependent protein kinase, catalytic subunit
EdU: 5-Ethynyl-2′-deoxyuridine
ER: Estrogen Receptor
GO: Gene Ontology
Her2: Human Epidermal growth factor Receptor 2
HR: Homologous Recombination
LDH: Lactate Dehydrogenase
MAP: Microtubule Associated Protein
MN: Micronuclei
MNC: Multi-Nucleated Cells
NBUD: Nuclear Budding
NHEJ: Non-Homologous End Joining
NPB: Nucleoplasmic Bridges
PARP: Poly ADP Ribose Polymerase
PI: Propidium Iodide
PR: Progesterone Receptor
SAC: Spindle Assembly Checkpoint
SEM: Standard Error of Mean
Rb: Retinoblastoma protein
TCGA: The Cancer Genome Atlas
TNBC: Triple Negative Breast Cancer

## Notes

### Competing Interest Statement

The authors have declared no competing interest.

