## Supplemental Figures for "Targeting LDHC dysregulates the cell cycle and improves sensitivity to cisplatin and olaparib"

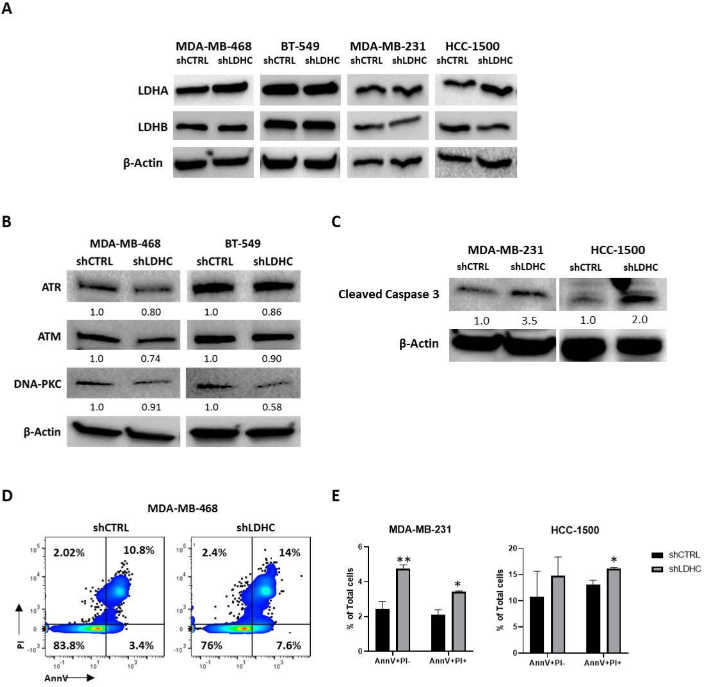


**Supplementary Figure 1.** ***LDHC*** **silencing does not alter LDHA/LDHB expression, and increases DNA damage and apoptosis.** A, Western blot of LDHA and LDHB, with β-actin as loading control. B, Western blot of DNA damage repair pathway mediators ATR and ATM (homologous recombination) and DNA-PKC (non-homologous end joining). Numbers under each lane represent mean densitometry values (arbitrary units) normalized to β-actin from three independent experiments. C, Western blot of cleaved caspase 3 in MDA-MB-231 and HCC-1500 shCTRL and shLDHC cells. Numbers under each lane represent mean densitometry values (arbitrary units) normalized to β-actin from three independent experiments. D, Representative AnnexinV/PI flow cytometry plots of MDA-MB-468 shCTRL and shLDHC cells. E, Annexin V/PI flow cytometric quantification of MDA-MB-231 and HCC-1500 shCTRL and shLDHC cells. Statistical analysis comparing shCTRL vs shLDHC performed using Student's t-test. Error bars represent standard error of mean (±SEM) from three independent replicates. *$p \leq 0.05$, ** $p \leq 0.01$.

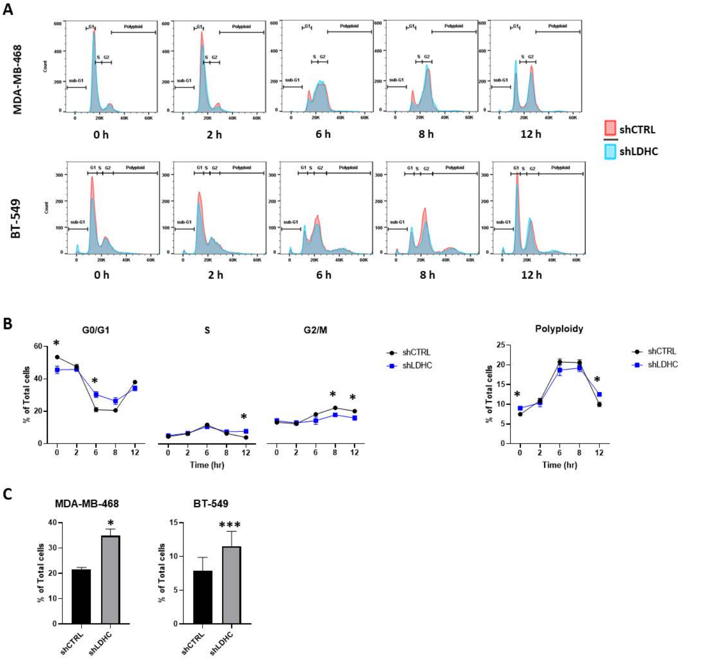


**Supplementary Figure 2. Aberrant cell cycle upon *LDHC* silencing.** A, Representative time-course of cell cycle histograms in synchronized MDA-MB-468 and BT-549 cells using PI flow cytometry. B, (Left) Time-course of cell cycle distribution of synchronized BT-549 cells using PI flow cytometry (error bars represent ± standard deviation). (Right) Frequency of BT-549 cells with polyploidy over a time-course of 12 h. C, Quantification of sub-G1 cell population at 12 h post thymidine block release. Statistical analysis comparing shCTRL vs shLDHC performed using Student's t-test. Error bars represent standard error of mean (±SEM) from three independent replicates. *$p \leq 0.05$, *** $p \leq 0.001$

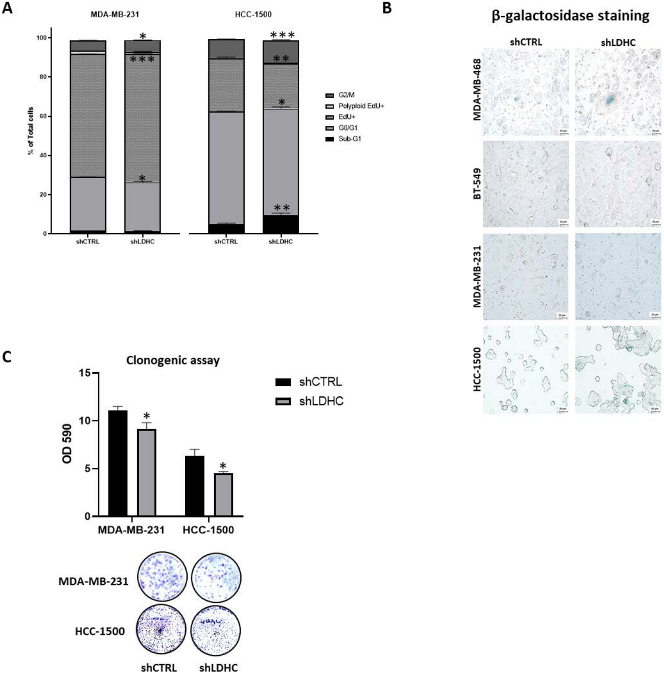


**Supplementary Figure 3. Aberrant cell cycle and cell fate upon *LDHC* silencing.** A, Representative asynchronous cell cycle profile of MDA-MB-231 and HCC-1500 cells, determined by EdU/PI flow cytometry. B, Visualization of senescence, determined by β-galactosidase staining. C, Crystal violet quantification at OD590 and representative image of MDA-MB-231 and HCC-1500 clonogenic assay. Statistical analysis comparing shCTRL vs shLDHC performed using Student's t-test. Error bars represent standard error of mean (±SEM) from three independent replicates. *$p \leq 0.05$, ** $p \leq 0.01$, *** $p \leq 0.001$

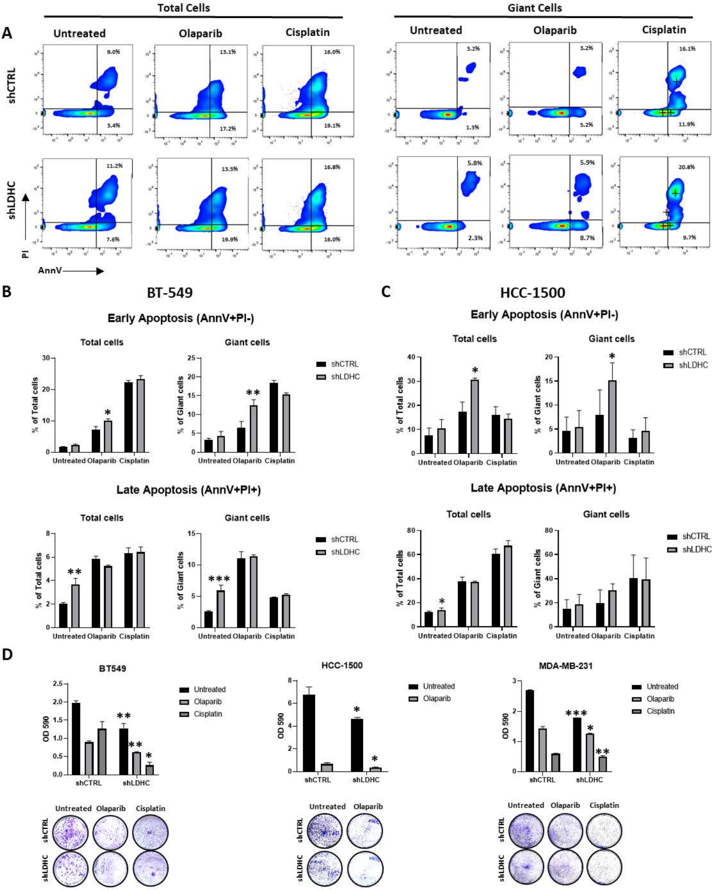


**Supplementary Figure 4. *LDHC* silencing improves sensitivity of breast cancer cells to DNA damage inducers and DNA damage repair inhibitors.** A, Representative Annexin V/PI flow cytometry plots of MDA-MB-468 cells after 72hrs of treatment with olaparib or cisplatin. B, Annexin V/PI flow cytometric quantification of BT-549 cells after 72hrs of treatment with olaparib or cisplatin. C, Annexin V/PI flow cytometric quantification of HCC-1500 cells after 72hrs of treatment with olaparib or cisplatin. D, BT-549, HCC-1500 and MDA-MB-231 clonogenic assay at 14 days of culture post-treatment (72h). Cisplatin treatment of HCC-1500 completely abolished clonogenic ability of shCTRL and shLDHC cells and hence is not depicted in the figure. Statistical analysis comparing shCTRL vs shLDHC performed using Student's t-test. Error bars represent standard error of mean (±SEM) from three independent replicates. *p≤0.05, **p≤0.01, ***p≤0.001

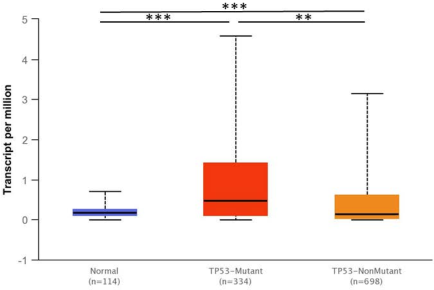


**Supplementary Figure 5. *LDHC* expression in *TP53* mutant breast cancer.** Box plot of *LDHC* mRNA expression in normal breast and TP53 mutant and non-mutant (wild-type) tumors using the TCGA Breast Cancer dataset. Data was retrieved and visualized using the UALCAN web-portal (available at http://ualcan.path.uab.edu). $^{**}p \leq 0.01$, $^{***}p \leq 0.001$
