## Supplemental Tables for "Targeting LDHC dysregulates the cell cycle and improves sensitivity to cisplatin and olaparib"

**Supplementary Table 1. List of antibodies**

| Antibody | Catalog no. | Manufacturer | Host | Concentration |
|---|---|---|---|---|
| Acetylated alpha-tubulin (Lys40) | 5335 | Cell Signaling | Rabbit | 1:200 (IF) |
| Alpha-tubulin | ab52866 | Abcam | Rabbit | 1:1000 (WB), 1:200 (IF) |
| ATM (D2E2) | 2873 | Cell Signaling | Rabbit | 1:1000 (WB) |
| ATR (E1S3S) | 13934 | Cell Signaling | Rabbit | 1:1000 (WB) |
| Aurora A | 14475 | Cell Signaling | Rabbit | 1:1000 (WB) |
| Beta-Actin | 4970 | Cell Signaling | Rabbit | 1:1000 (WB) |
| Beta-tubulin | 2128 | Cell Signaling | Rabbit | 1:1000 (WB), 1:200 (IF) |
| BubR1 | ab28193 | Abcam | Sheep | 1:500 (WB), 1:100 (IF) |
| Caspase 3 | 9662 | Cell Signaling | Rabbit | 1:1000 (WB) |
| Cdc25C | 4688 | Cell Signaling | Rabbit | 1:1000 (WB) |
| CDK6 | 3136 | Cell Signaling | Mouse | 1:1000 (WB) |
| CKS2 | ab155078 | Abcam | Rabbit | 1:1000 (WB) |
| Cyclin B1 | 12231 | Cell Signaling | Rabbit | 1:1000 (WB) |
| Cyclin D1 | 2978 | Cell Signaling | Rabbit | 1:1000 (WB) |
| Cyclin E2 | 4132 | Cell Signaling | Rabbit | 1:1000 (WB) |
| DNA-PKcs (E6U3A) | 38168 | Cell Signaling | Rabbit | 1:1000 (WB) |
| LDHC | ab52747 | Abcam | Rabbit | 1:500 (WB) |
| Mad2L1 | ab97777 | Abcam | Rabbit | 1:1000 (WB) |
| MAP1B | ab11266 | Abcam | Mouse | 1:200 (IF) |
| Myt1 | 4282 | Cell Signaling | Rabbit | 1:1000 (WB) |
| P16 | 80772 | Cell Signaling | Rabbit | 1:1000 (WB) |
| P18 | 2896 | Cell Signaling | Mouse | 1:1000 (WB) |
| P21 | 2947 | Cell Signaling | Rabbit | 1:1000 (WB) |

| | | | | |
|---|---|---|---|---|
| P27 | 3686 | Cell Signaling | Rabbit | 1:1000 (WB) |
| P53 | 2527 | Cell Signaling | Rabbit | 1:1000 (WB) |
| Phalloidin-Alexa Fluor 568 | A12380 | Thermo Fisher | | 1:100 (IF) |
| Phospho- gamma H2AX (S139) | ab11174 | Abcam | Rabbit | 1:1000 (WB), 1:200 (IF) |
| Phospho-cdc2 (Tyr15) | 4539 | Cell Signaling | Rabbit | 1:1000 (WB) |
| Phospho-Cdc25C (Ser216) | 4901 | Cell Signaling | Rabbit | 1:1000 (WB) |
| Phospho-Chk1 (Ser345) | 2348 | Cell Signaling | Rabbit | 1:1000 (WB) |
| Phospho-Chk2 (Thr68) | 2197 | Cell Signaling | Rabbit | 1:1000 (WB) |
| Phospho-Myt1 (Ser83) | 4281 | Cell Signaling | Rabbit | 1:1000 (WB) |
| Phospho-Wee1 (Ser642) | 4910 | Cell Signaling | Rabbit | 1:1000 (WB) |
| Wee1 | 13084 | Cell Signaling | Rabbit | 1:1000 (WB) |

WB, western blotting; IF, immunofluorescence

**Supplementary Table 2. Differentially expressed cell cycle genes.** Differentially expressed genes after LDHC silencing in MDA-MB-468 cells as determined by the Cell Cycle RT2 Profiler qPCR array.

| Gene Symbol | Fold Regulation |
|---|---|
| AURKA | 4.60 |
| AURKB | 2.80 |
| BIRC5 | 4.28 |
| BRCA1 | 2.24 |
| BRCA2 | 2.83 |
| CCNA2 | 4.03 |
| CCNB1 | 3.27 |
| CCNB2 | 2.42 |
| CCND3 | 2.77 |
| CCNF | 4.15 |
| CDC20 | 3.29 |
| CDC25A | 2.90 |
| CDC25C | 3.50 |
| CDC6 | 2.32 |
| CDK1 | 3.40 |
| CDK2 | 2.29 |
| CDKN3 | 2.21 |
| CHEK2 | 2.39 |
| CKS2 | 2.05 |
| E2F1 | 2.51 |
| GTSE1 | 3.58 |
| KPNA2 | 2.36 |
| MAD2L1 | 2.50 |
| MKI67 | 4.12 |
| RAD51 | 2.24 |
| STMN1 | 2.06 |

**Supplementary Table 3. Summary of observed characteristics in LDHC-silenced breast cell lines**

| | **MDA-MB-468** | **BT-549** | **MDA-MB-231** | **HCC-1500** |
|---|---|---|---|---|
| | ***Parental cell properties*** | | | |
| Molecular subtype | Basal-like | Basal-like | Basal-like | Luminal A |
| BRCA | Wild-type | Wild-type | Wild-type | Wild-type |
| p53 | mutant | mutant | mutant | negative |
| Rb | negative | negative | positive | positive |
| | *LDHC silencing phenotypes* | | | |
| Giant cells | Increase | Increase | Increase | Increase |
| Polyploidy | Increase | Increase | **low baseline level, no change** | **low baseline level, no change** |
| DNA damage | Increase | Increase | Increase | Increase |
| Apoptosis | Increase | Increase | Increase | Increase |
| Long-term survival/Clonogenicity | Decrease | Decrease | Decrease | Decrease |
| Microtubule instability | Increase | Increase | Increase | Increase |
| Cell population in G1 | Decrease | Decrease | Decrease | Decrease |
| Cell population in G2/M | Increase | Increase | Increase | Increase |
| Mitotic slippage | Increase | Increase | **ND** | **ND** |
| Cell cycle arrest | Increase | Increase | **ND** | **ND** |
| Senescence | No change | No change | No change | **Increase** |
| Sensitivity to Cisplatin | Increase | Increase | Increase | **No change** |
| Sensitivity to Olaparib | Increase | Increase | Increase | Increase |
